# Behavioral Paradigm for the Evaluation of Stimulation-Evoked Somatosensory Perception Thresholds in Rats

**DOI:** 10.1101/2023.05.04.537848

**Authors:** Thomas J. Smith, Yupeng Wu, Claire Cheon, Arlin A. Khan, Hari Srinivasan, Jeffrey R. Capadona, Stuart F. Cogan, Joseph J. Pancrazio, Crystal T. Engineer, Ana G. Hernandez-Reynoso

## Abstract

Intracortical microstimulation (ICMS) of the somatosensory cortex via penetrating microelectrode arrays (MEAs) can evoke cutaneous and proprioceptive sensations for restoration of perception in individuals with spinal cord injuries. However, ICMS current amplitudes needed to evoke these sensory percepts tend to change over time following implantation. Animal models have been used to investigate the mechanisms by which these changes occur and aid in the development of new engineering strategies to mitigate such changes. Non-human primates are commonly the animal of choice for investigating ICMS, but ethical concerns exist regarding their use. Rodents are a preferred animal model due to their availability, affordability, and ease of handling, but there are limited choices of behavioral tasks for investigating ICMS. In this study, we investigated the application of an innovative behavioral go/no-go paradigm capable of estimating ICMS-evoked sensory perception thresholds in freely moving rats. We divided animals into two groups, one receiving ICMS and a control group receiving auditory tones. Then, we trained the animals to nose-poke – a well-established behavioral task for rats – following either a suprathreshold ICMS current-controlled pulse train or frequency-controlled auditory tone. Animals received a sugar pellet reward when nose-poking correctly. When nose-poking incorrectly, animals received a mild air puff. After animals became proficient in this task, as defined by accuracy, precision, and other performance metrics, they continued to the next phase for perception threshold detection, where we varied the ICMS amplitude using a modified staircase method. Finally, we used non-linear regression to estimate perception thresholds.

Results indicated that our behavioral protocol could estimate ICMS perception thresholds based on ∼95% accuracy of rat nose-poke responses to the conditioned stimulus. This behavioral paradigm provides a robust methodology for evaluating stimulation-evoked somatosensory percepts in rats comparable to the evaluation of auditory percepts. In future studies, this validated methodology can be used to study the performance of novel MEA device technologies on ICMS-evoked perception threshold stability using freely moving rats or to investigate information processing principles in neural circuits related to sensory perception discrimination.

## Introduction

Intracortical microstimulation (ICMS) of the somatosensory cortex via microelectrode arrays (MEAs) has been successfully used to evoke cutaneous and proprioceptive sensations in amputees and individuals with spinal cord injuries (Armenta Salas et al., 2018; Bjånes et al., 2022; Christie et al., 2022; Page et al., 2021). These sensations can provide somatosensory feedback for closed-loop brain-machine interfaces and neuroprosthetics (Carè et al., 2022), which has been demonstrated to improve the control of robotic arms (Flesher et al., 2021). However, once implanted into the brain, achieving long-term stability of perception thresholds with these devices has been challenging (Callier et al., 2015; Hughes et al., 2021; Urdaneta et al., 2022) due to multifactorial failure of the interface. These failures include surpassing the safety limits of electrical microstimulation (Kramer et al., 2019; Pancrazio et al., 2017; Shannon, 1992), foreign body response that can isolate the MEAs from the surrounding neural tissue (Rajan et al., 2015), neuroinflammation that leads to neuronal loss (Ereifej et al., 2018; Potter et al., 2012), and material cracking and delamination (Barrese et al., 2013). Despite the promises of using ICMS to restore sensation, these failure modes pose a barrier for more widespread use. Because of this, research to improve the long-term reliability of ICMS is needed. The majority of pre-clinical studies investigating ICMS involve non-human primates; however, ethical concerns and costs limit their use (Bailey & Taylor, 2016; Carvalho et al., 2019; Pankevich, 2012). Rodents have been widely used to investigate the recording performance of MEAs due to their availability, affordability, and ease of handling (El-Ayache & Galligan, 2020; A. S. Koivuniemi et al., 2011). However, the use of this model organism for evaluating ICMS-induced somatosensory perceptions has been hindered by the limited behavioral paradigms available for this purpose.

To our knowledge, three behavioral paradigms have been described in the literature for assessing ICMS in the primary somatosensory cortex of rodents (A. Koivuniemi et al., 2011; Lycke et al., 2023; Öztürk et al., 2019; Urdaneta et al., 2021). These behavioral tasks use either a freely moving passive avoidance psychophysical detection task, a freely moving active avoidance conditioning paradigm, or a head-fixed go/no-go task. All were successful at detecting thresholds for up to 33 weeks with 70-95% accuracy; however, all three paradigms involve water deprivation for up to 36 hours prior to behavioral testing (A. Koivuniemi et al., 2011; Öztürk et al., 2019) which can produce stress (Vasilev et al., 2021) and confound chronic assessments. Alternative behavioral paradigms that use food-restriction have been described for the testing of auditory thresholds. An example of this is the well-established nose-poke behavioral paradigm (Abolafia et al., 2011; Riley et al., 2021; Schindler et al., 1993), a behavioral paradigm where a food-deprived rat is introduced into an operant conditioning chamber and trained to nose-poke through a hole on a side wall upon presentation of an auditory tone followed by a sugar pellet reward. While this behavioral task has been shown to be highly accurate with ∼90% discrimination accuracy scores (Riley et al., 2021; Sloan et al., 2009) and effective for auditory psychophysical testing, it has not been used to assess ICMS-induced somatosensory perceptions because no adaptations of the task have been made to suit this need.

Here we describe an innovative operant conditioning behavioral task to effectively assess ICMS-evoked sensory perception thresholds. We adapted the well-established and validated nose-poke auditory task into a food positive reinforcement go/no-go behavioral paradigm in food-deprived, freely moving rats with a mild passive avoidance positive-punishment air-puff. We implanted MEAs into Sprague-Dawley rats, targeting the forelimb area of the left primary somatosensory cortex (S1FL) and delivered electrical stimulation to modulate the neural activity and evoke artificial sensory percepts. We compared the accuracy of this task for ICMS perception thresholds with the accuracy of auditory tone discrimination for validation of the novel behavioral paradigm. Our results show that this behavioral protocol could estimate ICMS perception thresholds based on ∼95% accuracy of all rat nose-poke responses to the conditioned stimulus, validating its use for future ICMS perception threshold investigations.

## Material and Methods Ethics Statement

All animal handling, housing and procedures were approved by The University of Texas at Dallas IACUC (protocol #21-15) and in accordance with ARRIVE guidelines.

### Animal Use

We used six (N=6) male Sprague-Dawley rats (Charles River Laboratories Inc., Houston, TX, US) that were single-housed in standard home cages under a reverse 12-hour day/night cycle. We food-deprived the animals four consecutive days per week to a 90% free-feeding level that was redefined weekly to promote consistent performance during the behavioral task (Schindler et al., 1993) and given ad libitum access to food three consecutive days per week. Their weight was recorded on the last day of the week with ad libitum access to food, and before every behavioral session during the four consecutive days of food deprivation to assess welfare of the animal. If the weight before the behavioral session was below 90% of its recorded control weight, we provided supplemental rodent feed pellets to provide additional nourishment and excluded the animal from behavioral experimentation until the 90% free-feeding control weight was restored. Animals were given dustless reward pellets (F0021, Bio-Serv, Flemington, NJ, US) as positive reinforcement for the behavioral paradigm. These pellets contain a balanced caloric profile enriched with amino acids, carbohydrates, fatty acids, vitamin, and mineral mix to ensure the nutritional wellbeing of the animals despite food deprivation. In addition, we provided rats with supplemental regular food pellets (5LL2 - Prolab® RMH 1800, LabDiet, St. Louis, MO, US) after each behavioral session to maintain weight. This supplemental feed was calculated based on the number of reward pellets eaten during each behavioral session. Animals had ad libitum access to water at all times while in their standard home cages.

Rats were randomized and divided into two groups. The first was the experimental group, which received implantation with a multi-shank MEA (MEA-PI-A3-00-12-0.01-[1-2]-3-0.25-0.25-1-1SS; Microprobes for Life Science, Gaithersburg, MD, US) consisting of 12 Pt/Ir (70% Pt, 30% Ir, 0.01 MΩ) microwires of 75 μm diameter, insulated with polyamide. The tips of each microwire had an exposed geometric surface area ranging between 6000 and 9000 μm^2^. The MEA design has two rows of six microwires each, which slant in opposing directions ranging in length between 0.5 - 2 mm (Figure 1A). Each MEA includes an additional 2 mm microwire that serves as the reference electrode. The experimental group received ICMS (n=3) during the behavioral task. The second group was a control group (n=3), which underwent a sham surgery and received auditory tones during the behavioral task. The sham surgery consisted of a craniotomy and durotomy procedure comparable with the experimental group without implantation of the MEA. The goal of the control group was to compare the accuracy of the behavioral paradigm presented here. The operant chamber apparatus was thoroughly cleaned with a 70% ethanol solution between each session to help eliminate any distracting scents between animal subjects. After completing the behavioral testing, the animals in the ICMS group were subjected to the same behavioral task without electrical stimulation. This was done to act as an intragroup negative control to validate ICMS as the only interpreted conditioning cue by verifying changes in accuracy during the absence of a stimulus.

**Figure 1.**
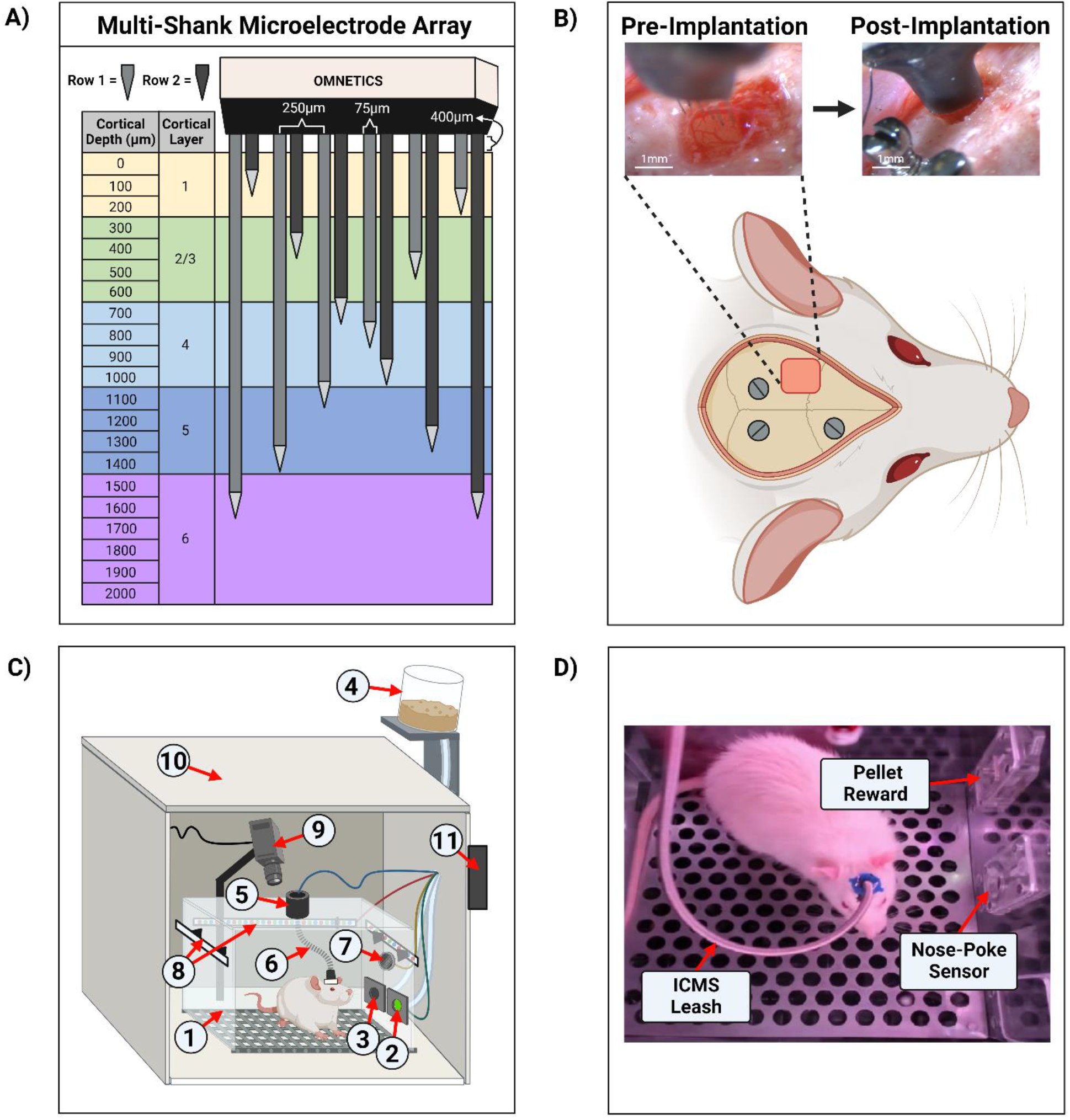
Experimental setup and microelectrode array implantation. **(A)** Diagram of the twelve-shank MEA with opposing slanted rows penetrating all layers of the somatosensory cortex **(B)** Example of an implantation surgery (craniotomy, durotomy, and microelectrode array insertion) within the left primary somatosensory cortex, forelimb area (S1FL). Three stainless steel screws were inserted into the skull for ground/counter electrodes and headcap anchors. **(C)** Illustration of the operant conditioning chamber setup used for animal behavior. The setup contains: (1) operant conditioning chamber, (2) nose-poke sensor hole, (3) sugar pellet reward hole, (4) pellet dispenser, (5) commutator, (6) ICMS leash, (7) speaker, (8) RGB LED strips, (9) webcam/camera, (10) noise reduction chamber, (11) microcontroller board hub. **(D)** Screenshot from a behavioral live stream session depicting a real-world view. In the image, the sugar pellet reward hole, nose-poke sensor hole, and the ICMS leash were shown.

### Surgical Procedure

Rats underwent a surgical procedure for sham and MEA implantation as previously described (Sturgill et al., 2022). Briefly, animals were anesthetized using vaporized isoflurane (1.8-2.5%) mixture with medical grade oxygen (500 mL/min; SomnoSuite® for Mice & Rats, Kent Scientific Corporation, Torrington, CT, US). The surgical team monitored vital signs throughout the surgical procedure while body temperature was maintained using a controlled far-infrared warming pad (PhysioSuite® for Mice & Rats, Kent Scientific Corporation, Torrington, CT, US). The scalp was shaved and animals were mounted onto a digital stereotaxic frame (David Kopf Instruments, Tujunga, CA, US). The skin at the surgical site was cleaned using three alternating applications of betadine and alcohol wipes. A subcutaneous injection of 0.5% bupivacaine hydrochloride (Marcaine, Hospira, Lake Forest, IL, US) was given at the intended incision site. An incision was made through the midline of the scalp, muscles, and connective tissue. Next, the skull was leveled and centered in the stereotaxic frame using bregma, lambda, and the sagittal suture as references (± 0.1 mm). Three holes were then drilled into the skull to insert stainless-steel bone screws (Stoelting Co., Wood Dale, IL, USA) (Figure 1B). Then, a 2 mm × 3 mm craniotomy was made targeting the S1FL (AP: -0.5 mm, ML: 4 mm), followed by a durotomy (Figure 1B). The surgeon secured the ground wire to one of the mounted bone screws and implanted the MEA to a cortical depth of ∼1.6 mm using a precision-controlled inserter (NeuralGlider, Actuated Medical, Inc., Ann Arbor, MI, US) (Figure 1B). Implantation within the cranial window was done to avoid disruption of major surface blood vessels (He et al., 2022; Kozai et al., 2010). The implant site was then sealed with a biocompatible, transparent silicone elastomer adhesive (Kwik-Sil, World Precision Instruments, Sarasota, FL, US), followed by a dental cement head cap to tether the MEA to the skull while also reducing the likelihood of contamination and infection. Then, the incision was closed using surgical staples and tissue adhesive (GLUture, World Precision Instruments, Sarasota, FL, US). At the end of the surgical procedure, we injected each animal with 0.05 mL/kg intramuscular cefazolin (Med-Vet International, Mettawa, IL, US) for antibiotic prophylaxis together with topical application triple-antibiotic ointment around the incision site. For analgesia, we administered either 0.15 mL/kg of subcutaneous slow-release (Buprenorphine SR-LAB, ZooPharm, LLC., Laramie, WY, US) or 0.5 mL/kg of extended-release (Ethiqa XR, Fidelis Animal Health, North Brunswick, NJ, US) buprenorphine depending on availability of the substance. When necessary, we administered a dose of buprenorphine after 72 hours post-surgery if the animal showed signs of pain. Lastly, we provided sulfamethoxazole and trimethoprim oral suspension (200 mg/40 mg/5 mL, Aurobindo Pharma, Dayton, NJ, US) in the animals’ drinking water (1 mL/100 mL drinking water) as an additional antibiotic for 7 days post-surgery.

### Behavioral Operant Chamber, Equipment and Software

Figure 1C&D illustrates the behavioral operant chamber used for this study. The go/no-go behavioral paradigm was conducted within a commercially available operant conditioning chamber (OmniTrak, Vulintus, Inc., Lafayette, CO, US). This chamber had two holes in one of the side walls, one containing an infrared break-beam sensor (nose-poke sensor) and a second hole connected to a precision pellet dispenser. In addition, the nose-poke hole had the capability of delivering a mild air-puff from a medical-grade compressed air cylinder tube as positive punishment. This air-puff was controlled via a pneumatic solenoid (SKUSKD1384729, AOMAG) connected to an Inland Nano microcontroller through a relay switch to deliver air to the nose-poke sensor hole. A rotating commutator (76-SR-12, NTE Electronics, Bloomfield, NJ, US) was bolted at the top of the operant chamber to allow the animals to roam free while connected to an external stimulator (PlexStim, Plexon Inc., Dallas, TX, US) for ICMS. A custom cord was designed to connect the animal to the commutator for ICMS, incorporating an Omnetics (A79021-001, Omnetics Connector Corporation, Minneapolis, MN, US) adapter and surrounded with a stainless-steel spring cable shielding (#6Y000123101F, Protech International Inc., Boerne, TX, US) to protect the wires against biting. For the auditory control group, auditory tones were presented through a mini speaker (Product ID: 3923, Adafruit Industries, New York City, NY, US) that was placed inside the chamber and connected to a PC’s headphone auxiliary port. The chamber was illuminated via an RGB LED strip controlled by the Inland Nano microcontroller. A webcam (960-001105, Logitech, Lausanne, CH, US) was mounted to the chamber to record a live video stream of the animal during behavioral sessions. Finally, the chamber was enclosed inside a sound-reduction chamber equipped with a fan for cooling and air circulation. All modules were connected and controlled by an ATMEGA2560 microcontroller board hub (OmniTrak Controller V3.0, Vulintus Inc., Lafayette, CO, US), interfaced using custom MATLAB (R2022b, Mathworks, Natick, MA, US) software. The RBG LED strip and solenoid valve required a supplemental 12V 2A DC power supply to power the devices.

In addition, we developed a custom MATLAB GUI application (Supplementary Figure 1) that simultaneously controls and displays the behavioral task parameters, monitors animal performance, and records session data. While a behavioral session is active, the application feeds the session video live stream from the operant chamber to the researcher, as shown in Supplementary Figure 1. Furthermore, this GUI included specialized buttons for the researcher to annotate instances during each session where we deemed the animals distracted (e.g., grooming or turning away from the sensors/modules for the entire trial duration) for exclusion from analysis. After each session, a second researcher validated the annotations offline to reduce bias. Additional features of the GUI application include a button for manually dispensing sugar pellets, the ability to record voltage transients throughout the session, and the capability to choose which electrode channels are delivered ICMS. This custom MATLAB and UI/UX behavior software is available as an open-source package on GitHub (https://github.com/Neuronal-Networks-and-Interfaces-Lab/Stimulation-Evoked_Perception_Behavioral_Software.git).

### Electrical Stimulation and Auditory Parameters

Electrical stimulation for ICMS was delivered to 10 electrode sites simultaneously per implanted MEA. The stimulation parameters selected for this work were previously established by another group and validated to evoke somatosensory percepts in rats (Urdaneta et al., 2021). We used current-controlled, charge-balanced symmetric biphasic waveforms with a cathodal-leading phase, a frequency of 320 Hz, pulse width of 200 μs per phase, 40 μs interphase interval, with a 650 ms train duration (PlexStim, Plexon Inc., Dallas, TX, US). Current amplitudes used in this work ranged from 0-25 μA corresponding to a charge of 0-5 nC/ph. The maximum charge limit set for all experiments was 5 nC/ph per electrode stimulated simultaneously across ten channels. Seven to twelve days after implantation but before operant conditioning training, we estimated a provisional ICMS naïve perception threshold for each animal by slowly increasing the charge/phase across all 10 individually pulsed channels simultaneously from 0 to 5 nC/ph until a physical response (e.g., paw withdrawal) was observed. Once this provisional perception threshold was determined, we confirmed that the physical response was driven primarily by somatosensory ICMS and not motor activation by presenting the stimulus at the same charge intensity while the animal was anesthetized (1.8-2.0% isoflurane). This naïve perception threshold was subsequently used as the starting known threshold for the go/no-go behavioral paradigm. Voltage transients during ICMS were recorded by connecting the external stimulator to an oscilloscope (TBS1052B, Tektronix, Beaverton, OR, US).

For the auditory control group, auditory tone parameters were derived from prior go/no-go paradigms (Engineer et al., 2008; Green et al., 1979; Sloan et al., 2009). In our experiment, we used a carrier frequency of 6 kHz pure tone sinusoidal wave with a 100 kHz sampling rate, 500 ms tone duration, and a 50 ms beginning/end tone ramp duration. Using a sound level meter (Extech Instruments, Nashua, NH, US), the produced output intensity of this auditory training tone was measured to be ∼90 dB in reference to the sound pressure level (SPL) of 0 dB, which is the intensity of sound waves relative to the minimum threshold of human hearing.

### Go/No-Go Behavioral Training

We trained rats on the go/no-go behavioral paradigm following a three-tier protocol. Namely, Shaping, Shape2Detect, and Detection, as shown in Figure 2. Each tier is designed to gradually train every animal to nose-poke following a presented stimulus (ICMS or auditory tone) to receive a reward pellet in the go/no-go paradigm as shown in Figure 3A. Before training began, animals were habituated for a minimum of 10 hours until the animal tolerated handling and head restraint for at least two consecutive minutes. This habituation allowed for manipulation of the animals and connection of the implanted MEA to the rotating commutator hardware before each behavioral session. During the habituation period, the animals were fed reward pellets to incentivize the reward-seeking behavior.

**Figure 2.**
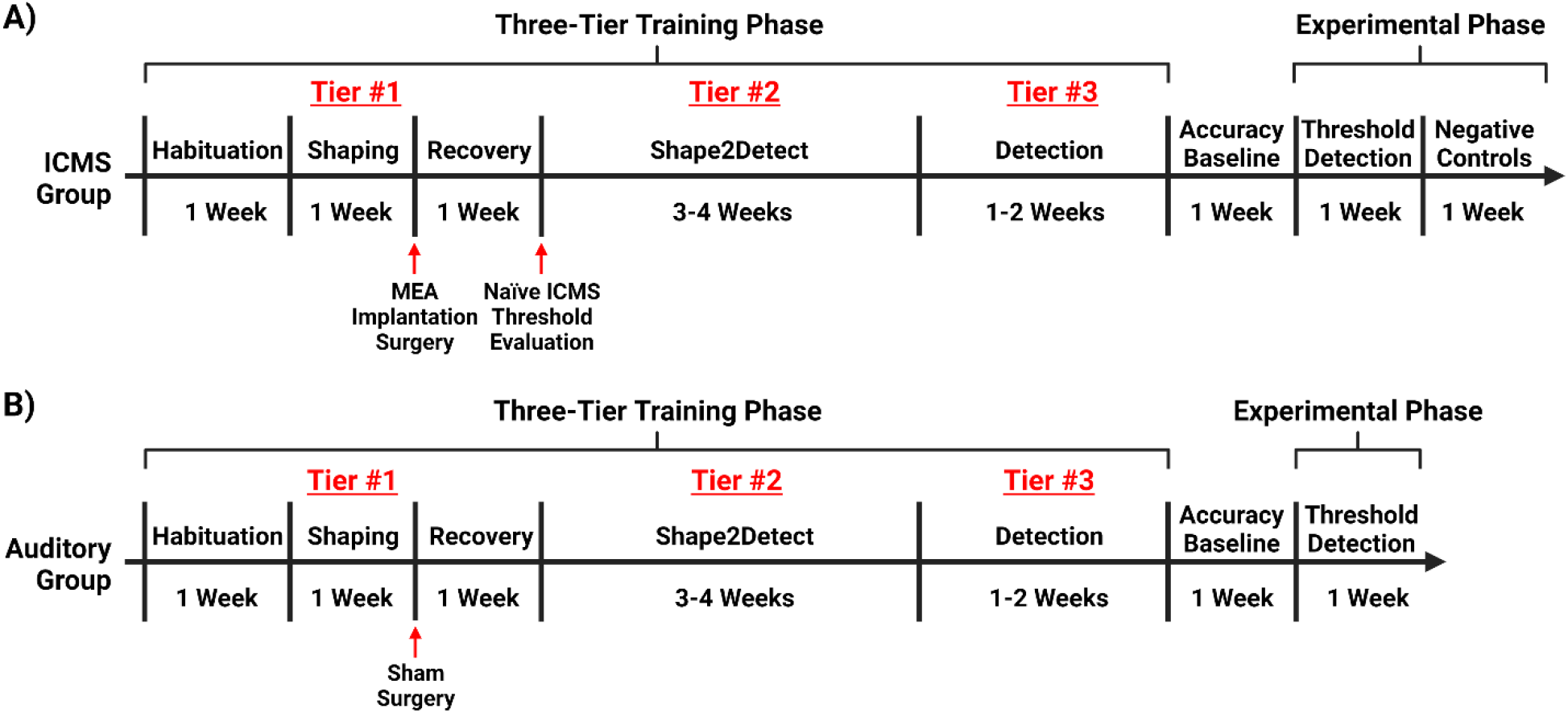
Experimental timeline. Timeline for training rats on the go/no-go behavioral paradigm. **(A)** Training for rats in the ICMS experimental group with an extended phase where no ICMS is presented, acting as an intragroup negative control. **(B)** Training for rats in the auditory control.

**Figure 3.**
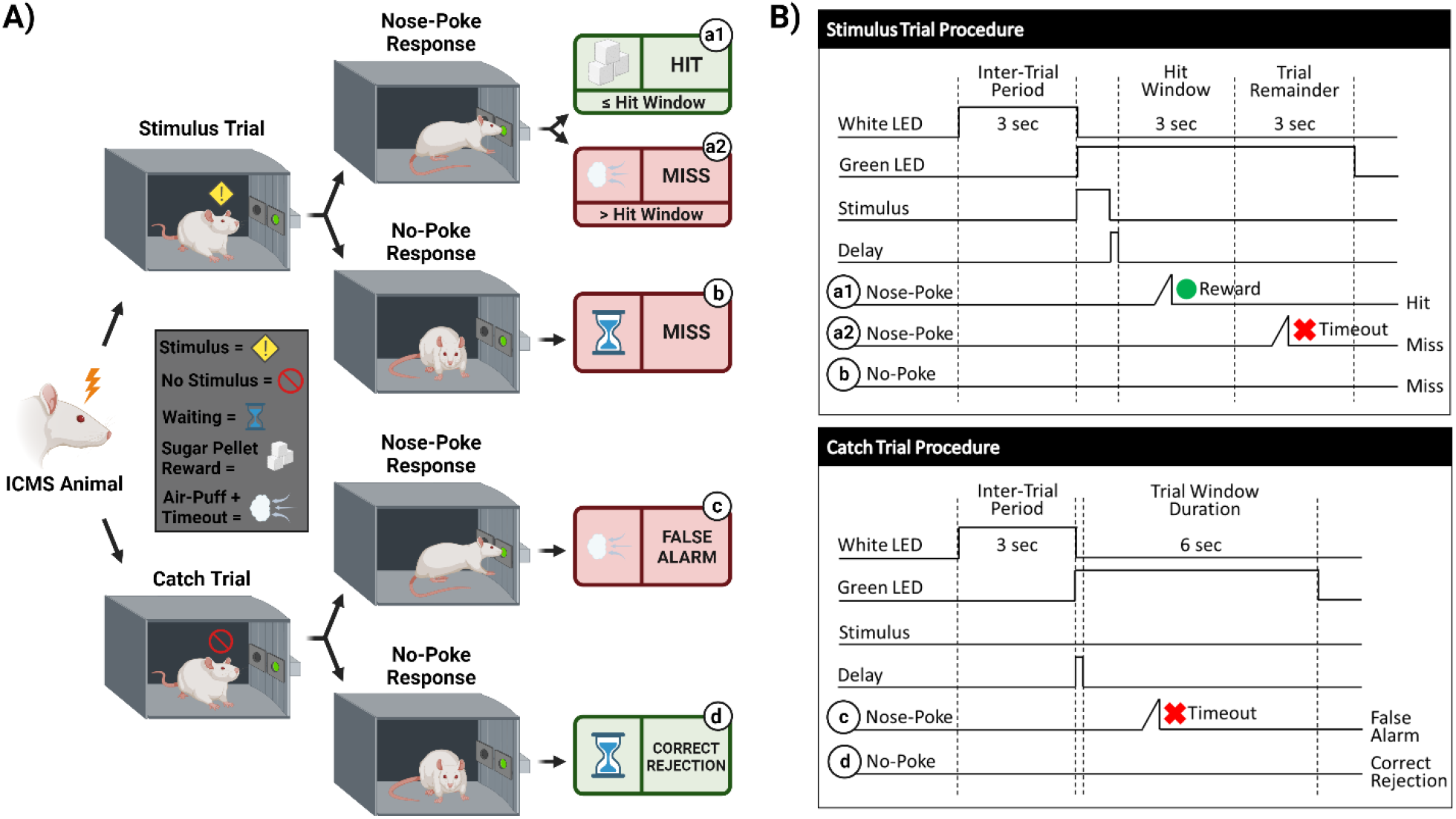
Behavioral paradigm for go/no-go task. **(A)** Visualization of the go/no-go behavioral paradigm with possible responses to ICMS. **(B)** Illustration of the go/no-go behavioral paradigm outlining trial types. Schematic shows differences between the stimulus trials (top) and the catch trials (bottom). Depending on the response to the presented trial type, the animal can either receive a sugar pellet reward (hit) symbolized by the green circle, an 8 s timeout sequence + air puff (false alarm) symbolized by the red x, or nothing (miss/correct rejection). A 150 ms delay immediately following a stimulus presentation is used, where the nose-poke sensor does not trigger.

### First Tier: Shaping

Shaping was the first tier for the go/no-go training, which consisted of one-hour sessions, five days per week. The goal of this phase was to train the animal on the nose-poke behavior task via positive reinforcement. First, the animal was introduced into the operant chamber and allowed to freely roam. The chamber was illuminated with white light via the RGB LED strip. After three seconds, the RGB LED strip was configured to illuminate with green light for an indefinite amount of time, indicating a trial had begun. A pellet reward was dispensed when the animal nose-poked through the nose-poke hole as a positive-reinforcement to promote this behavior unless the animal poked within the first 150 ms of the trial. This delay was incorporated to prevent accidental nose-pokes from occurring at the start of a trial. After the animal nose-poked, the green light turned back to white light for an inter-trial period of three seconds. If the animal nose-poked during the inter-trial period, no reward pellet was dispensed. When needed, we manually dispensed pellets when animals approached the nose-poke hole, even if the animal did not poke to encourage exploration. Animals were considered proficient in the Shaping task when they received 100+ reward pellets for two consecutive sessions without manual pellets dispensed. After passing this tier, they received either surgery for MEA implantation, or sham surgery. If a rat did not meet the 100+ pellet reward within 10 sessions, the animal was excluded from the study.

### Second Tier: Shape2Detect

Shape2Detect was the second tier for the go/no-go task training, as shown in Figure 2. During this phase, animals were trained to nose-poke only upon presentation of either the ICMS at their pre-established naïve threshold or the auditory training tone at ∼90 dB SPL, depending on their group allocation. We began each session by placing the animal into the apparatus once per day, four days per week for 60-minute-long sessions. At the start of the session, the operant chamber was illuminated by white light from the RGB LED strip. When each trial began, the RGB LED strip changed to green light to indicate the beginning of a trial (Figure 3B). During this phase, animals were presented with two types of trials: stimulus trials or catch trials as outlined in Figure 3A&B. A stimulus trial was defined as the presentation of the ICMS or auditory tone; whereas a catch trial consisted of an absence of stimulation or sound. Positive punishment was tied to the catch trial to reinforce the rat’s ability to ignore trials in the absence of stimulus and discourage nose-poking freely. Stimulus and catch trials were presented sequentially in trial windows followed by a 3 second inter-trial period of white light. The trial window duration varied as time progressed throughout the session, as shown in Table I. For the first 20 minutes, the trial window duration was set to 3 seconds. The next ten minutes had trial durations of 4 seconds, the following ten minutes durations of 5 seconds, and the final ten minutes durations of 6 seconds. Throughout the session, the likelihood of a stimulus trial being presented versus a catch trial was varied. The first ten minutes had an 83.3% probability of presenting a stimulus trial (with a 16.7% probability of catch trials) and then changed until the last ten minutes had a 50% probability of presenting a stimulus trial (50% probability of catch trials). The rationale for varying this probability was to increase the frequency of stimulus exposure at the beginning of the session, providing the animal ample opportunities to associate the stimulus presentation with a reward. Then, we decreased the frequency of the stimulus exposure as the session progressed to avert continuous poking and encourage discriminatory decision making. Finally, the hit window and timeouts were also varied throughout the session (see Table I). The hit window was defined as the duration of time after the presentation of a stimulus during which the animal can nose-poke and receive a pellet reward (Figure 3B). A hit was determined if an animal nose-poked during this hit window. If an animal nose-poked after the hit window (trial remainder) or during a catch trial, it received a mild-air puff as a punishment and triggered a timeout period, characterized by red light illumination. The first instance was classified as a miss for quantification purposes; the latter as a false alarm. If the animal poked during the timeout period, it received an air-puff and additional time was added to the timeout. The pressure of the air-puff was adjusted as needed so that it was enough to prevent timeouts but not to completely deter the animal from nose-poking. Furthermore, if the animal failed to nose-poke for ten stimulus trials in a row, the session would be paused and resumed only after the animal nose-poked again. Finally, a correct rejection was defined as the animal refraining from nose-poking during a catch trial.

**Table I.** Shape2Detect behavioral training task parameters

| Session Time<br>(min) | Trial Window<br>Duration (s) | Stimulus Trial<br>Probability (%) | Hit Window<br>(s) | Timeout<br>(s) |
| --- | --- | --- | --- | --- |
| 0-9 | 3 | 83.3 | 3 | 2 |
| 10-19 | 3 | 71.4 | 3 | 3 |
| 20-29 | 4 | 66.7 | 4 | 3 |
| 30-39 | 5 | 60.0 | 5 | 5 |
| 40-60 | 6 | 50.0 | 3 | 8 |

In the context of this study, hits and correct rejections were considered true responses, whereas misses and false alarms were considered false responses. Animals were considered proficient in the Shape2Detect task if they met four conditions for two consecutive sessions: 1) at least a 75% accuracy (Equation 1), 2) 75% precision (Equation 2), 3) 75% hit rate (Equation 3) score, and 4) received at least 100 reward pellets.

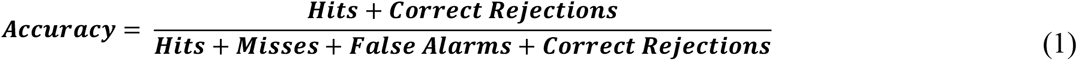

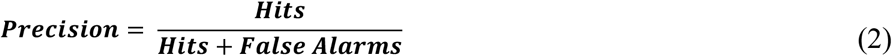

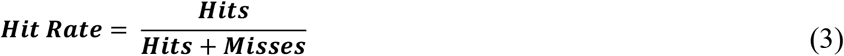

### Third Tier: Detection

Detection was the third tier for the go/no-go task training (Figure 2). The goal of this phase was to maximize animal accuracy during consistently paced trials with invariable parameters. This phase of training was similar to the Shape2Detect task but used fixed behavioral parameters throughout the 60-minute-long sessions. These parameters outlined in Figure 3B were the same as those used during the last 20 minutes of the Shape2Detect sessions (i.e., 6 second trial window duration, 3 second hit window, 50% probability of presenting a stimulus trial, and 8 second timeouts). Animals were considered proficient when they showed at least 75% accuracy, 75% precision, 75% hit rate, 75% correct rejection rate (Equation 4), and 75% F1-score (Equation 5) with at least a 1.5 d-prime (*d’*) score (Equation 6) in three total sessions. The F1-score is a measure of performance in binary classification that considers the harmonic mean, in this case, of an animal’s precision and hit rate scores. The *d’* metric is another performance indicator and common statistical measure used in psychophysical detection tasks and signal detection theory to quantify a subject’s ability to accurately distinguish between a signal and noise within a given task.

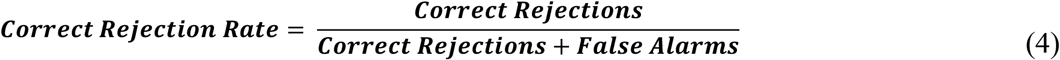

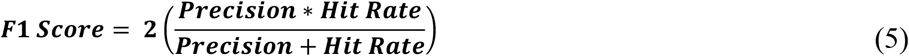

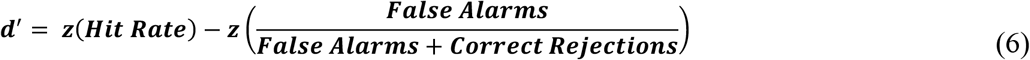

After the training on the go/no-go paradigm was completed, animals underwent five additional Detection sessions to assess baseline accuracy and subject consistency before proceeding to the go/no-go perception threshold detection task.

### Go/No-Go Perception Threshold Detection Task

After rats were fully trained in the go/no-go behavioral paradigm, they were introduced to a dynamic perception threshold detection task that implemented a modified version of the up/down staircase method (A. Koivuniemi et al., 2011; Levitt, 1971), as shown in Supplementary Figure 2. The goal of this task was to approximate an estimation of an animal’s perception threshold value. The first 20 minutes of every perception threshold detection task began with all ICMS stimulus trials presented at the naïve threshold intensity and with 50% probability (catch trials were presented as the alternative). For the remainder of the session, the naïve threshold intensity was presented with a 33.3% probability, while a dynamic charge intensity was also presented with 33.3% probability (the remainder probability presented a catch trial). The dynamic charge intensities were presented following the modified staircase method (Figure 4A). First, we presented the dynamic charge intensity value at the maximum naïve threshold intensity. If the rat perceived the dynamic charge intensity value and nose-poked, the dynamic charge intensity value was decreased by the step size variation outlined in Table II. If the rat did not nose-poke, the dynamic charge intensity value was increased. This up/down staircase methodology was followed throughout the session.

**Figure 4.**
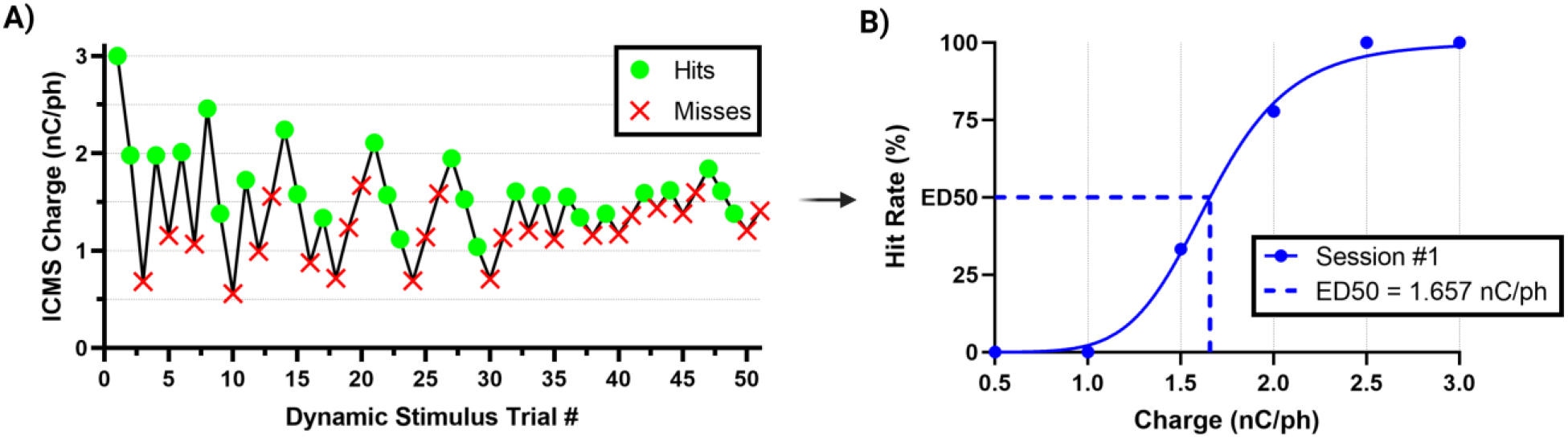
Estimation of ICMS perception thresholds. **(A)** Representative nose-poke response data from the modified staircase presentation of ICMS during a typical threshold detection session. **(B)** Representative quantal dose-response, non-linear regression plot showcasing transformed hit/miss animal response data into percent hit rate based on binned (ranges of 0.5 nC/ph pulsed across all individual channels simultaneously) charge amplitude values presented. Effective charge (dose) at 50% hit rate (ED50) were used to estimate the ICMS perception thresholds.

**Table II.** Dynamic stimulus step size variation throughout a one-hour session

| Session Time<br>(min) | Step size variation |  |
| --- | --- | --- |
|  | Charge Intensity<br>(nC/ph) | Tone Intensity<br>(% amplitude) |
| 0-19 | No variation | No variation |
| 20-29 | $1.00 \pm 0.40$ | $20.00 \pm 5.00$ |
| 30-39 | $0.60 \pm 0.20$ | $10.00 \pm 3.00$ |
| 40-49 | $0.40 \pm 0.10$ | $1.00 \pm 0.30$ |
| 50-60 | $0.20 \pm 0.05$ | $0.10 \pm 0.03$ |

For the auditory stimulus trials, dynamic tone intensity values were determined by modulating the sinusoidal wave amplitude of the training tone. Increases in sinusoidal wave amplitude resulted in a louder and more intensely perceived tone, while decreases produced a quieter and less intense tone. To create a scale for estimating auditory tone thresholds, the amplitude of the training tone was normalized to a range of 0-100%, where 0% represented silence (0 dB SPL) and 100% represented the maximum intensity of the training tone (∼90 dB SPL). Similar to the ICMS variation, initial trials in the perception threshold detection task were presented at the maximum training tone intensity of 100% amplitude with a 50% probability. The remaining trials followed the modified staircase method where changes in dynamic tone intensity values were presented to the rats based on their response behavior. Step size variations of auditory tone intensity in percent amplitude are outlined in Table II.

### Estimation of Threshold Perception

We estimated perception thresholds using non-linear regression (Equation 7) in a quantal dose-response non-linear regression (Liu et al., 2022; Müller & Schmitt, 1990) in the GraphPad Prism Software ([Agonist] vs. normalized response --Variable slope, Prism, v9.5.1). In Equation 7, *x* represents the linear dose in charge/phase or percent amplitude, *y* denotes the normalized response of the percent hit rate from 0-100%, and the Hillslope represents the slope factor or steepness of the curve shared globally between all perception threshold detection sessions per animal. We binned the dynamic stimulus trial values into increments of 0.5 nC/ph stimulated across all individual channels simultaneously for the ICMS group and 1% sinusoidal wave amplitude for the auditory group to establish a quantal response (Figure 4B). We defined the effective dose in charge/phase or percent amplitude needed to produce a 50% hit rate response (ED50) as previously demonstrated (Müller et al., 1990). In this equation, we constrained ED50 so that it must be greater than zero. Finally, perception threshold values were estimated individually for all animals in the ICMS and auditory groups, using the ED50 data collected across five go/no-go perception threshold detection task sessions.

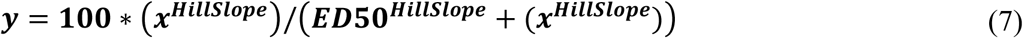

### Data Analysis and Statistics

All data analysis was conducted through custom MATLAB (R2022b) scripts, GraphPad Prism (v9.5.1, GraphPad Software, Boston, MA, US), or Statgraphics Centurion 19 (v19.4.04, Statgraphics Technologies, Inc., The Plains, VA, US). In MATLAB, we evaluated signal detection theory parameters (Macmillan & Creelman, 2005) for all behavioral sessions, including: accuracy, precision, hit rate, correct rejection rate, F1-score, and *d’* (equations 1-6). If a session contained either zero hits, misses, false alarms, or correct rejection responses – all of which are denominators in equations (1-6) – then their values were adjusted in order to prevent behavioral performance scores of infinities using a commonly accepted approach (Macmillan & Creelman, 2005). An arbitrary value of 0.5 was added to the metric that had a score of zero (e.g., hits, misses, false alarms, or correct rejections), meanwhile this arbitrary value of 0.5 was subtracted from its non-zero counterpart. For example, if a session contained 119 hits and zero misses, then the adjusted values would be 118.5 hits and 0.5 misses. Then, we generated confusion matrices based on these calculations for each group to highlight the overall accuracies, hit rates, and correct rejection rates during the accuracy baseline Detection task sessions.

GraphPad Prism was used to calculate the perception threshold values. Furthermore, we calculated the average training time for each group. For statistical analysis, unpaired two-sample t-tests were used to determine significant differences between the ICMS and auditory groups. We conducted a one-tailed paired sample t-test between the ICMS results and the intragroup negative control for further validation of this methodology. We analyzed tests of normality in the data using the Shapiro-Wilk test and confirmed results by examination of their respective QQ plots. Lastly, we performed an equivalence test using Statgraphics Centurion 19 to further investigate if the average ICMS group accuracy was statistically similar or different than the average auditory group accuracy. The upper and lower differential limits were determined from the 95% CI range of the difference between means (Hazra, 2017). All results are reported as the mean ± SEM. We defined statistical significance as p < 0.05.

## Results

All animals remained above the 90% weekly weight limit for the entire duration of this study, demonstrating that food restriction did not affect their weight. Furthermore, 70% of animals completed the study with at least a 20% increase in overall weight compared to their first shaping session; the remaining animals showed less than 5% weight loss (Supplementary Table I). All animals passed the Shaping task in less than 10 sessions, resulting in no exclusions from the study due to poor performance.

After implantation of the MEA into the S1FL for animals in the ICMS group, we proceeded with testing of the naïve threshold. All three animals showed a paw withdrawal in the right forepaw, corresponding to the contralateral implant location; two animals responded reliably at 3 nC/ph pulsed across all individual channels simultaneously, and one responded at 4 nC/ph. Voltage transients from each microelectrode array channel were recorded to confirm set stimulation parameters outlined within the Electrical Stimulation and Auditory Parameters subsection. Figure 5 displays a representative in-vivo current-controlled voltage transient of a single channel recorded during a 3 nC/ph pulse train.

**Figure 5.**
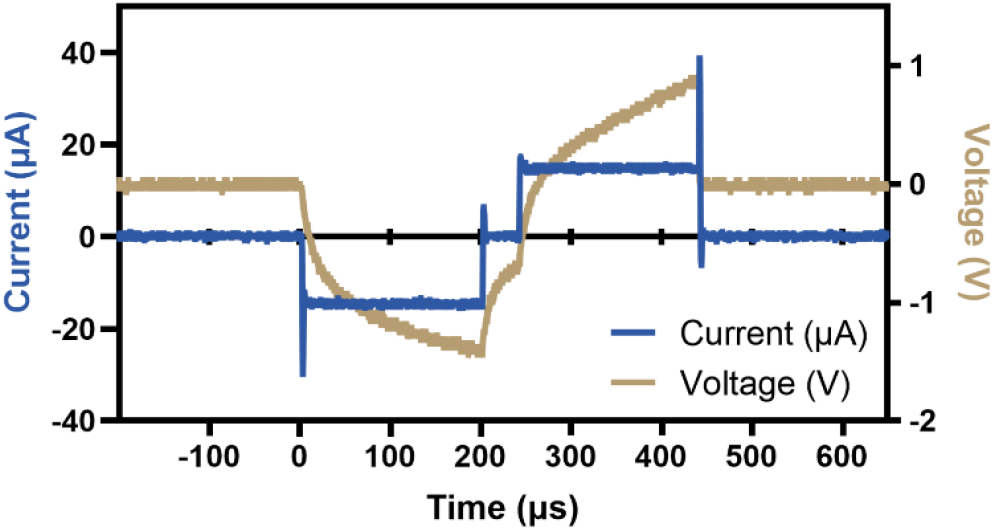
Representative voltage transient. Shown is a representative 3 nC/ph current controlled voltage transient used to stimulate each microelectrode array channel individually.

Voltage transients showed that the electrode delivered electrical stimulation consistently and remained unchanged throughout the sessions and validated that the applied current amplitude was delivered as set in the MATLAB custom GUI.

## Go/No-Go Behavioral Training

Figure 6 provides the assessment of behavioral proficiency in the go/no-go task. As shown in Figure 6A, animals in the ICMS group took an average of 15.3 ± 2.2 sessions in total between Shaping, Shaping2Detect and Detection tasks, while animals in the auditory group took an average of 20.7 ± 3.7 sessions (p=0.28). This number of sessions corresponds 4-5 weeks of training for the animal to become proficient in the go/no-go behavioral task.

**Figure 6.**
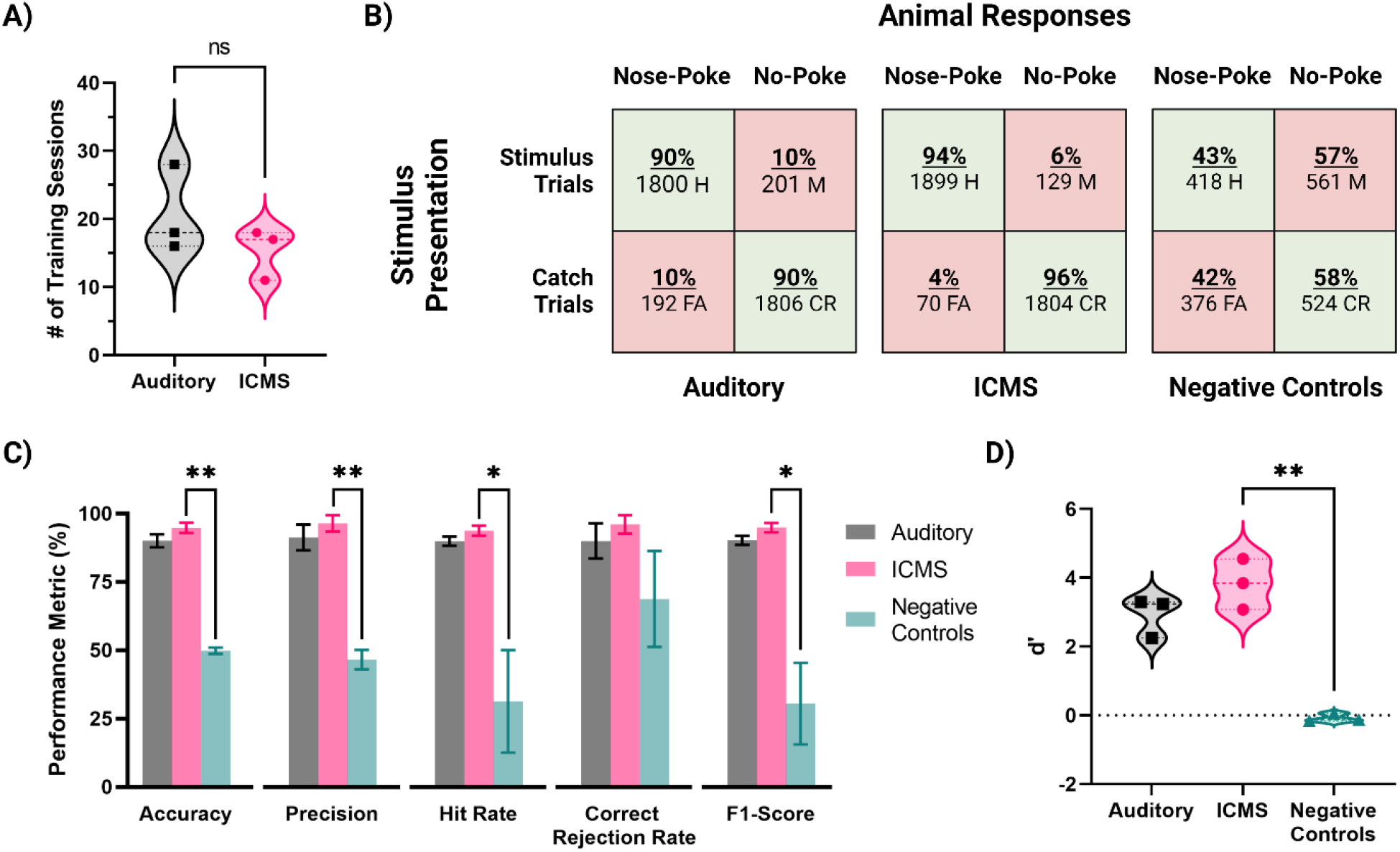
Behavioral performance metrics for the ICMS and auditory groups, and for negative control stimulation. **(A)** Training time for each group, in number of sessions needed to pass the training phase. **(B)** Confusion matrices showing presented trials (rows) and animal responses (columns). Values depict all animal response data from five baseline accuracy sessions. **(C)** Behavioral performance metrics, including accuracy, precision, hit rate, correct rejection rate, and F1-Score. Data are shown as mean ± SEM. **(D)** Average scores of the *d’* metric.

Then, we proceeded to assess the baseline performance on the Go/No-Go behavioral task of each animal in five post-training sessions. Figure 6B shows the overall distribution of the total presented trials (rows) and animal responses (columns) for each group, represented in the form of confusion matrices. There was a total of 3,902 trials presented for the ICMS animals, including stimulus (2,028) at the naïve threshold and catch (1,874) trials. In comparison, the auditory group received 3,999 total trials (stimulus trials: 2,001, catch trials: 1,998). Animals in both, auditory and ICMS groups showed similar hit rates (auditory = 90%, ICMS = 94%), showing that the animals are correctly poking upon most stimulation trials. Similarly, animals in both groups had a high correct rejection rate (auditory = 90%, ICMS = 96%). These results indicate that both groups of animals were able to greatly recognize a stimulus signal and respond with a nose-poke. In contrast, when the stimulation was turned off for the ICMS group (negative control) the hit rate dropped down to only 43% and correct rejections to only 58%, signifying random poking. Figure 6C outlines the accuracy performance metrics for all groups. The average accuracy scores between the ICMS (94.7 ± 1.9%) and auditory (90.0 ± 2.4%) groups were comparable to one another (p = 0.19). In addition, the equivalence test performed subsequently demonstrated that the accuracy for both groups was equivalent (p = 0.03). In contrast, the ICMS and negative controls (49.8 ± 1.2%) were significantly different (p=0.002). The average precision scores between the ICMS (96.4 ± 3.0%) and auditory (91.2 ± 4.7%) groups were comparable (p = 0.41); the difference between ICMS and negative controls (46.6 ± 3.6%) was statistically significant (p = 0.008). The average hit rates between the ICMS (93.7 ± 1.8%) and auditory (89.9 ± 1.7%) groups comparable (p = 0.19); differences between the ICMS group and negative controls (31.3 ± 18.8%) were statistically significant (p = 0.04). The average correct rejection rates between the ICMS (96.0 ± 3.3%) and auditory (89.9 ± 6.4%) groups were comparable (p = 0.45); difference between ICMS and negative controls (68.7 ± 17.6%) did not reach statistical significance (p = 0.10). These correct rejection rates show that all animals were able to identify catch trials regardless of stimuli type.

The average F1-scores between the ICMS (94.9 ± 1.7%) and auditory (90.2 ± 1.7%) groups were comparable (p = 0.12). The difference between the ICMS and negative controls (30.5 ± 14.9%) was statistically significant (p = 0.03), further demonstrating that animals are only poking upon stimulus presentation. In addition, the average d’ scores (Figure 6D) between the ICMS (3.82 ± 0.43) and auditory (2.93 ± 0.34) groups were comparable (p = 0.18); the difference between ICMS and negative controls (-0.08 ± 0.07) was found to be statistically significant (p = 0.008), demonstrating that the animals are able to distinguish between stimulus and catch trials.

### Estimated Perception Thresholds

Across five sessions of the go/no-go perception threshold detection task, we estimated the perception thresholds for all animals in the auditory and ICMS groups. Figure 7A (left) shows the estimated perception threshold values for individual sessions for each animal in the auditory group. The perception threshold between sessions for each animal showed a standard deviation from the mean ranging from 0.27 to 0.90% of the sinusoidal wave amplitude. Figure 7A (right) shows the summary statistics, where the perception threshold was estimated at 1.74 ± 0.19% sinusoidal wave amplitude. Figure 7B (left) shows the estimated perception threshold values for individual sessions for each animal. Animals in the ICMS group showed a small standard deviation from the mean ranging from 0.16 to 0.45 nC/ph pulsed across all individual channels simultaneously in the perception thresholds across all five sessions. Figure 7B (right) shows that the average perception threshold across all animals is 1.64 ± 0.15 nC/ph pulsed across all individual channels simultaneously.

**Figure 7.**
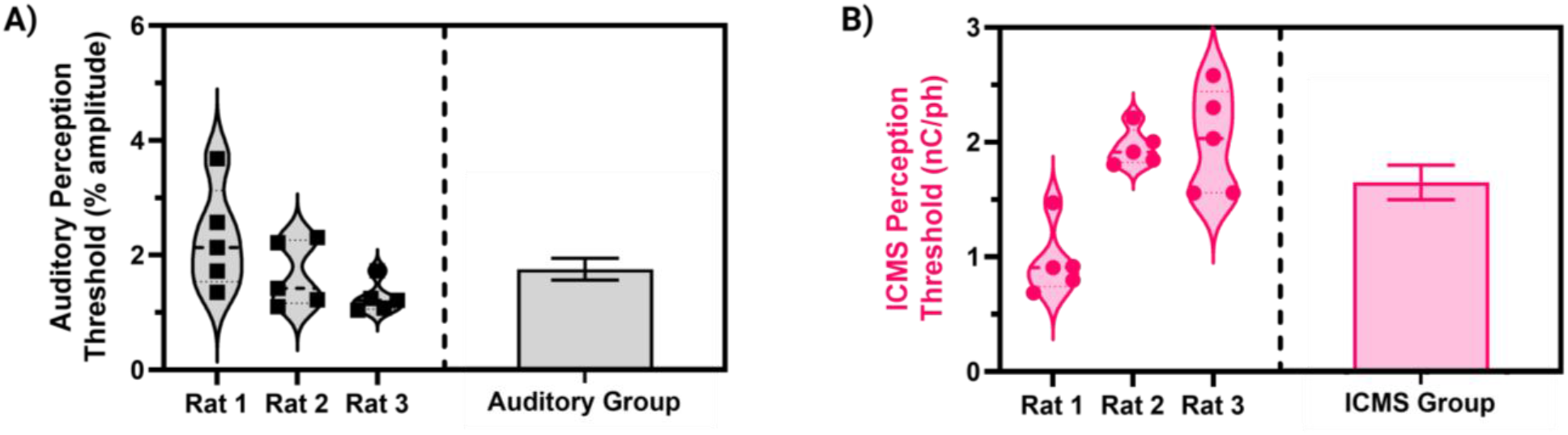
Estimated perception thresholds for the ICMS and auditory animal groups. **(A)** Estimated perception threshold values plotted for each auditory animal (left) and auditory group estimations (right) shown as mean ± SEM. **(B)** Estimated perception threshold values plotted for each ICMS animal (left) and ICMS group estimations (right) shown as mean ± SEM.

## Discussion

In this study, we developed and validated an innovative non-pain aversive, go/no-go behavioral paradigm based on a nose-poking task to quantify rat sensory perception thresholds in response to ICMS. Our results showed that this nose-poking paradigm could reliably assess stimulation-evoked sensory percepts in rats originating from ICMS in the S1FL and its accuracy was comparable to the well-established auditory discrimination task.

The study of auditory tone discrimination tasks in animals has a long and rich history in neuroscience research. Early studies in the 1970s focused on fundamental aspects of auditory perception in rats, such as their ability to detect pure tones and discriminate between tones of different frequencies and intensities (Kelly & Masterton, 1977). These studies laid the foundation for more complex auditory tasks developed in the following decades (Hui et al., 2009; Sloan et al., 2009). One such task is the go/no-go task, which once involved training rats to press a lever in response to a specific tone (the “go” tone) and withhold their response to other tones (“no-go” tones) (Engineer et al., 2008). Then, this go/no-go task was modified from lever-pressing to nose-poking because it was found to require less experimenter intervention for a naïve rat to reliably perform the task with the addition of a higher baseline rate of responding and lower between-group variability (Mekarski, 1988; Schindler et al., 1993). This nose-poke go/no-go behavioral paradigm has been used by multiple research groups and is widely accepted because of its straightforwardness to train rats with nose-poking being an innate exploration behavior, the hardware is available off-the-shelf and does not require complex motors and controls, and it has shown high accuracy rates of up to ∼90% (Riley et al., 2021; Sloan et al., 2009). Overall, the history of auditory tone discrimination tasks in rats highlights their broad utility as a model system for studying auditory perception and processing. For the development of the behavioral paradigm presented here, we built upon this nose-poke-based, go/no-go paradigm.

To validate the presented behavioral paradigm, we compared the ICMS group to an auditory discrimination control group. Using the auditory discrimination group as positive controls allowed us to establish an effective baseline to compare accuracy and reliability of our behavioral paradigm. Within our study, the auditory control group showed an accuracy of ∼90% and demonstrated an auditory tone threshold of approximately 2% amplitude (∼65 dB SPL), which is comparable to previous literature (Engineer et al., 2008; Riley et al., 2021; Sloan et al., 2009). These results validate our implementation of the nose-poke behavioral paradigm, and our method of using non-linear regression for estimating threshold perception. The ICMS group had a comparable accuracy to the auditory control group of ∼95%, which validates the use of this go/no-go nose-poke task for the assessment of ICMS perception. Furthermore, animals in the ICMS group underwent a negative control phase at the end of the study to confirm that the nose-poking behavior was neither random nor were the animals nose-poking on any confounding cues. Results from this second phase of the investigation yielded a 50% accuracy, which is an indication of random poking, which is consistent with the present methodology.

Using the validated quantal non-linear regression at the ED50 level, we established that the average electrical perception threshold across three animals was approximately 1.64 nC/ph pulsed across all 10 individual channels simultaneously with the lowest animal averaging 0.96 nC/ph. Previous animal behavioral paradigms have been developed to study sensory and visual perception via ICMS, including rodents, cats, non-human primates, and humans (Fernández et al., 2021; Lycke et al., 2023; Ni & Maunsell, 2010; Rousche & Normann, 1999; Tehovnik, 1996), which have identified different thresholds of perception. Urdaneta et al. (2022) demonstrated perception thresholds ranging between 6.4 and 10.7 nC/ph for rat cortex, when stimulating Ir electrode sites individually. The same group has demonstrated that delivering electrical stimulation through two or more electrode sites simultaneously can reduce the perception threshold (Kunigk et al., 2022) by at least 53% of the single site perception threshold. Other studies have shown lower perception thresholds using traditional microelectrode arrays in cat somatosensory cortex (Rousche & Normann, 1999) with an approximate threshold of 1.5 nC/ph; non-human primates between 1-2 nC/ph (Callier et al., 2015; Ferroni et al., 2017; Ni & Maunsell, 2010); and human studies ranging from 0.4-3 nC/ph (Fernández et al., 2021; Flesher et al., 2016; Hughes et al., 2021; Schmidt et al., 1996). A different study targeting the primary somatosensory cortex in mice (Lycke et al., 2023) found the lowest perception threshold of 0.25 nC/ph stimulating individual and multiple electrode sites simultaneously. It should be noted that stimulation parameters, MEAs, implantation targets, and number of electrode sites pulsed are not consistent between these studies. Nevertheless, results from these prior studies demonstrate broad consistency with the estimated perception thresholds in the present work.

Some Institutional Animal Care and Use Committees (IACUCs) require ad libitum access to water for a minimum of 1 hour for at least every 12 hours, which may further limit the deployment of previous water-restrictive behavioral paradigms to other research groups. Food restriction is preferred over water restriction by most IACUCs. In this paradigm we mildly restricted food intake, an approach ethically preferred over water deprivation, to ensure rodent engagement during the behavioral task. At the end of each session, animals were given supplemental feed to ensure appropriate nutrition. However, both water deprivation and food restriction have been associated with a stress response characterized by an upregulation of adrenal corticosterone (Dietze et al., 2016; Vasilev et al., 2021). It is unknown whether this stress response may play a role in the reliability of intracortical MEAs and stability of ICMS. Future work may consider methods to avoid food restriction while participating in the nose-poke task.

A final limitation of this study was the training time, resulting from having a mostly positive reinforcement behavioral task. Animals in this study underwent one week of Shaping, three to four weeks of Shape2Detect, one to two weeks of Detection and one week of the accuracy baseline Detection task assessment for a total of six to eight weeks of training. During this time, we could not assess perception thresholds, meaning that we could not assess changes during the first six to eight weeks post-implantation. Previous studies (Urdaneta et al., 2022) have reported training phases of up to eight weeks post implantation, comparable to the number of sessions required for training in the present paradigm. However, this acute phase is known for presenting changes to the MEA surrounding tissues, including myelin degeneration and glial encapsulation. Assessment during the acute phase would provide information regarding perception threshold and documented tissue response. In future studies, we will optimize the training time to assess perception thresholds as early as possible after implantation by increasing the probability of presenting a stimulus trial during the Shape2Detect and Detection phases of training and lowering the threshold to pass from one training stage to the next.

Despite these limitations, this study presents an effective behavioral paradigm for evaluating ICMS-evoked somatosensory percepts in rats. However, there are still known challenges associated with rat ICMS studies apart from establishing a reliable perception threshold indicator. For example, it has been well-documented that perception thresholds change over time (Bjånes et al., 2022; Callier et al., 2015; Hughes et al., 2021; A. Koivuniemi et al., 2011; Kunigk et al., 2022; Lycke et al., 2023). In the future we will employ this behavioral paradigm to study ICMS-evoked perception threshold stability of novel MEA device technologies that aim at improving the long-term reliability of the neural interface. Finally, the control software that we have developed for this paradigm is open-source and available to download at no cost. This will allow research groups who are interested in evaluating long-term stability of novel stimulating MEAs (especially those whose IACUC prefer food restriction over water deprivation in rodents) to easily adopt this go/no-go behavioral paradigm using hardware available off-the-shelf.

## Conclusion

In this study we presented a new, highly accurate behavioral paradigm to assess ICMS-evoked somatosensory perception thresholds. This paradigm builds upon well-established and accepted auditory discrimination tasks with comparable results, validating the go/no-go behavioral task for assessment of ICMS-evoked percepts. Full deployment of this paradigm establishes a new platform for elucidating the information processing principles in the neural circuits related to neuroprosthetic sensory perception and for studying the performance of novel MEA device technologies using freely moving rats. Future studies will assess how MEA design and cortical circuitry impacts stimulus response-time circuitry, threshold sensitivity, and selectivity discrimination for the primary somatosensory cortex.

## Data Availability Statement

The data is available upon request to the corresponding author. MATLAB custom GUI behavior software is available as an open-source package on GitHub (https://github.com/Neuronal-Networks-and-Interfaces-Lab/Stimulation-Evoked_Perception_Behavioral_Software.git).

## Ethics Statement

All aspects of this study were conducted under our 21-15 protocol endorsed by the Institutional Animal Care and Use Committee (IACUC) at the University of Texas at Dallas and were in accordance with the ARRIVE essential 10 guidelines.

## Author Contributions

**Thomas Smith:** Conceptualization, Methodology, Software, Validation, Formal Analysis, Investigation, Resources, Writing – Original Draft, Visualization, Project Administration. **Yupeng Wu:** Investigation. **Claire Cheon:** Investigation. **Arlin Khan:** Investigation. **Hari Srinivasan:** Investigation. **Jeffrey Capadona:** Conceptualization, Writing – Review and Editing, Funding Acquisition. **Stuart Cogan:** Conceptualization, Resources, Writing – Review and Editing, Funding Acquisition. **Joseph Pancrazio:** Conceptualization, Resources, Writing – Review and Editing, Project Administration, Funding Acquisition. **Crystal Engineer:** Methodology, Resources, Writing – Review and Editing. **Ana Hernandez-Reynoso:** Conceptualization, Methodology, Software, Formal Analysis, Writing – Review and Editing, Project Administration.

## Acknowledgements

The authors thank undergraduate Ian Okidhain for his contribution to the custom software and electrical circuitry used to develop the behavioral paradigm and apparatus setup. In addition, the authors thank undergraduate students Mihai Bendea, Fareeha Faruk, Mehak Kaul, Shreya Tirumala Kumara, Teresa Thai, Sophia Vargas, and Rebeca Villafranca for their contribution and assistance with the voltage transient and behavioral data collection. Lastly, the authors thank Alan Carroll for developing the Shape2Detect task used in this study.

---

**Supplementary Figure 1.**
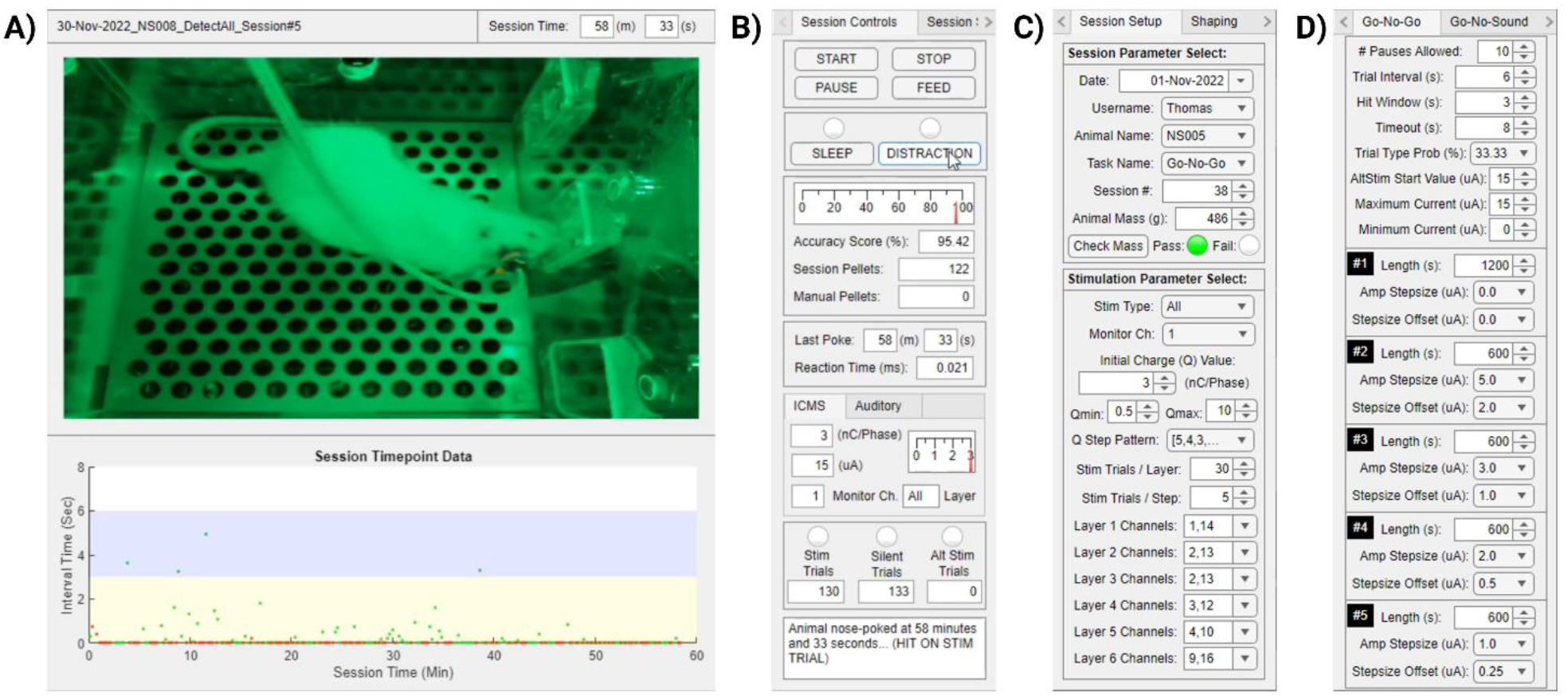
Custom MATLAB GUI application. **(A)** Main screen showing live video feed (top) from behavioral chamber during a stimulation trial, and nose-poke response data (bottom) throughout the session. **(B)** Session controls showing buttons to start, pause, and stop the session manually, and to manually feed a reward pellet. In addition, there are two buttons to mark times during the session when the animal is distracted or sleeping. The remainder of the panel displays live performance and behavioral task metrics such as session accuracy, number of reward pellets eaten, timepoint of last nose-poke, trial reaction time, stimulus intensity values, and a text box that presents various status updates. **(C)** Session setup panel displaying options for selecting the date, researcher, animal, task name, current session number and a button to confirm animal mass at or above 90% free feeding level. It also contains an ICMS parameter selection panel used to define the intensity of the stimulus and electrode channels used. **(D)** Example behavioral task panel showing the options for changing the go/no-go task parameters outlined in the study.

**Supplementary Figure 2.**
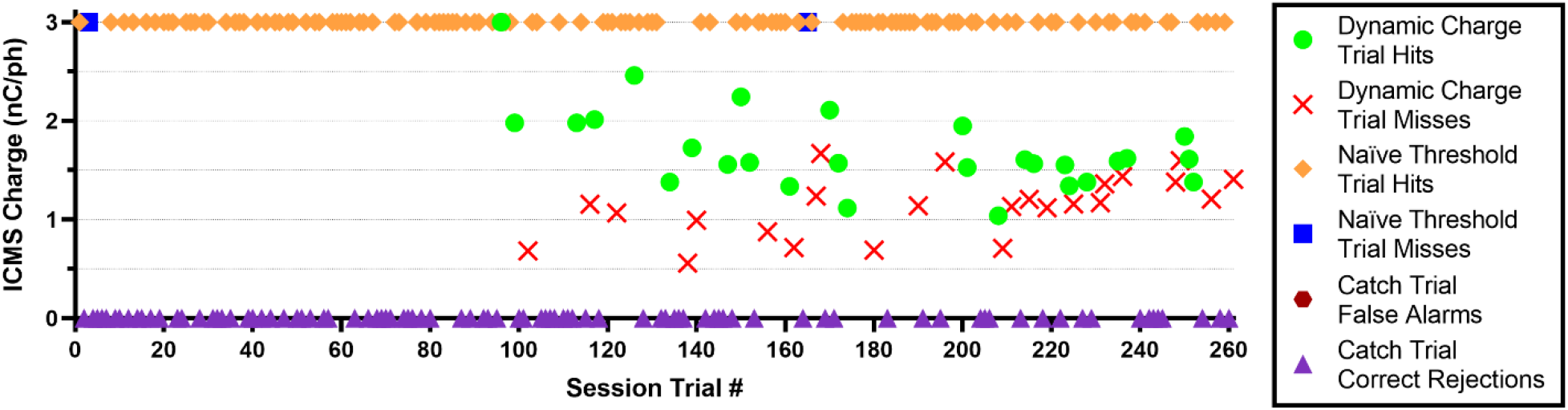
Perception Threshold Detection Task session showing all trial response types. **(A)** Representative animal response data from the go/no-go perception threshold detection task session. This chart plots an ICMS animal’s responses in hits or misses for the dynamic stimulus and naïve stimulus trials, and false alarms or correct rejections for the catch trials presented throughout a typical one-hour session. There were no catch trial false alarms present within this example session. Additionally, the dynamic charge trial hits and misses plotted here are equivalent to the hits and misses plotted in Figure 4A.

**Supplementary Table I.** Animal Weight Progression

| Animal Group | Animal | Session Weight |  |  |
| --- | --- | --- | --- | --- |
|  |  | First Session (g) | Last Session (g) | % Change |
| ICMS | Rat 1 | 412 | 578 | 40.29 |
| ICMS | Rat 2 | 355 | 517 | 45.63 |
| ICMS | Rat 3 | 522 | 503 | -3.64 |
| Auditory | Rat 1 | 452 | 569 | 25.88 |
| Auditory | Rat 2 | 550 | 523 | -4.91 |
| Auditory | Rat 3 | 362 | 436 | 20.44 |

## References

Abolafia, J., Martinez-Garcia, M., Deco, G., & Sanchez-Vives, M. (2011). Slow Modulation of Ongoing Discharge in the Auditory Cortex during an Interval-Discrimination Task. Frontiers in Integrative Neuroscience, 5. https://doi.org/10.3389/fnint.2011.00060

Armenta Salas, M., Bashford, L., Kellis, S., Jafari, M., Jo, H., Kramer, D., Shanfield, K., Pejsa, K., Lee, B., Liu, C. Y., & Andersen, R. A. (2018). Proprioceptive and cutaneous sensations in humans elicited by intracortical microstimulation. eLife, 7, e32904. https://doi.org/10.7554/eLife.32904

Bailey, J., & Taylor, K. (2016). Non-human Primates in Neuroscience Research: The Case against its Scientific Necessity. Alternatives to Laboratory Animals, 44(1), 43–69. https://doi.org/10.1177/026119291604400101

Barrese, J. C., Rao, N., Paroo, K., Triebwasser, C., Vargas-Irwin, C., Franquemont, L., & Donoghue, J. P. (2013). Failure mode analysis of silicon-based intracortical microelectrode arrays in nonhuman primates. Journal of Neural Engineering, 10(6), 066014. https://doi.org/10.1088/1741-2560/10/6/066014

Bjånes, D. A., Bashford, L., Pejsa, K., Lee, B., Liu, C. Y., & Andersen, R. A. (2022). Multi-channel intra-cortical micro-stimulation yields quick reaction times and evokes natural somatosensations in a human participant. medRxiv, 2022.2008.2008.22278389. https://doi.org/10.1101/2022.08.08.22278389

Callier, T., Schluter, E. W., Tabot, G. A., Miller, L. E., Tenore, F. V., & Bensmaia, S. J. (2015). Long-term stability of sensitivity to intracortical microstimulation of somatosensory cortex. Journal of Neural Engineering, 12(5). https://doi.org/10.1088/1741-2560/12/5/056010

Carè, M., Averna, A., Barban, F., Semprini, M., De Michieli, L., Nudo, R. J., Guggenmos, D. J., & Chiappalone, M. (2022). The impact of closed-loop intracortical stimulation on neural activity in brain-injured, anesthetized animals. Bioelectronic Medicine, 8(1), 4. https://doi.org/10.1186/s42234-022-00086-y

Carvalho, C., Gaspar, A., Knight, A., & Vicente, L. (2019). Ethical and Scientific Pitfalls Concerning Laboratory Research with Non-Human Primates, and Possible Solutions. Animals, 9(1). https://doi.org/10.3390/ani9010012

Christie, B., Osborn, L. E., McMullen, D. P., Pawar, A. S., Thomas, T. M., Bensmaia, S. J., Celnik, P. A., Fifer, M. S., & Tenore, F. V. (2022). Perceived timing of cutaneous vibration and intracortical microstimulation of human somatosensory cortex. Brain Stimulation: Basic, Translational, and Clinical Research in Neuromodulation, 15(3), 881–888. https://doi.org/10.1016/j.brs.2022.05.015

Dietze, S., Lees, K. R., Fink, H., Brosda, J., & Voigt, J.-P. (2016). Food Deprivation, Body Weight Loss and Anxiety-Related Behavior in Rats. Animals, 6(1).

El-Ayache, N., & Galligan, J. J. (2020). Chapter 28 - The Rat in Neuroscience Research. In M. A. Suckow, F. C. Hankenson, R. P. Wilson, & P. L. Foley (Eds.), The Laboratory Rat (Third Edition) (pp. 1003–1022). Academic Press. https://doi.org/https://doi.org/10.1016/B978-0-12-814338-4.00028-3

Engineer, C. T., Perez, C. A., Chen, Y. T. H., Carraway, R. S., Reed, A. C., Shetake, J. A., Jakkamsetti, V., Chang, K. Q., & Kilgard, M. P. (2008). Cortical activity patterns predict speech discrimination ability. Nature Neuroscience, 11(5). https://doi.org/10.1038/nn.2109

Ereifej, E. S., Rial, G. M., Hermann, J. K., Smith, C. S., Meade, S. M., Rayyan, J. M., Chen, K., Feng, H., & Capadona, J. R. (2018). Implantation of Neural Probes in the Brain Elicits Oxidative Stress [Original Research]. Frontiers in Bioengineering and Biotechnology, 6. https://doi.org/10.3389/fbioe.2018.00009

Fernández, E., Alfaro, A., Soto-Sánchez, C., Gonzalez-Lopez, P., Lozano, A. M., Peña, S., Grima, M. D., Rodil, A., Gómez, B., Chen, X., Roelfsema, P. R., Rolston, J. D., Davis, T. S., & Normann, R. A. (2021). Visual percepts evoked with an intracortical 96-channel microelectrode array inserted in human occipital cortex. The Journal of Clinical Investigation, 131(23). https://doi.org/10.1172/JCI151331

Ferroni, C. G., Maranesi, M., Livi, A., Lanzilotto, M., & Bonini, L. (2017). Comparative performance of linear multielectrode probes and single-tip electrodes for intracortical microstimulation and single-neuron recording in macaque monkey. Frontiers in Systems Neuroscience, 11. https://doi.org/10.3389/fnsys.2017.00084

Flesher, S. N., Collinger, J. L., Foldes, S. T., Weiss, J. M., Downey, J. E., Tyler-Kabara, E. C., Bensmaia, S. J., Schwartz, A. B., Boninger, M. L., & Gaunt, R. A. (2016). Intracortical microstimulation of human somatosensory cortex. Science Translational Medicine, 8(361), 361ra141–361ra141. https://doi.org/doi:10.1126/scitranslmed.aaf8083

Flesher, S. N., Downey, J. E., Weiss, J. M., Hughes, C. L., Herrera, A. J., Tyler-Kabara, E. C., Boninger, M. L., Collinger, J. L., & Gaunt, R. A. (2021). A brain-computer interface that evokes tactile sensations improves robotic arm control. Science, 372(6544), 831–836. https://doi.org/10.1126/science.abd0380

Green, M., Terman, M., & Terman, J. S. (1979). Comparison of yes-no and latency measures of auditory intensity discrimination. J Exp Anal Behav, 32(3), 363–372. https://doi.org/10.1901/jeab.1979.32-363

Hazra, A. (2017). Using the confidence interval confidently. J Thorac Dis, 9(10), 4125–4130. https://doi.org/10.21037/jtd.2017.09.14

He, F., Sun, Y., Jin, Y., Yin, R., Zhu, H., Rathore, H., Xie, C., & Luan, L. (2022). Longitudinal neural and vascular recovery following ultraflexible neural electrode implantation in aged mice. Biomaterials, 291, 121905–121905. https://doi.org/https://doi.org/10.1016/j.biomaterials.2022.121905

Hughes, C. L., Flesher, S. N., Weiss, J. M., Downey, J. E., Boninger, M., Collinger, J. L., & Gaunt, R. A. (2021). Neural stimulation and recording performance in human sensorimotor cortex over 1500 days. Journal of Neural Engineering, 18(4). https://doi.org/10.1088/1741-2552/ac18ad

Hui, G. K., Wong, K. L., Chavez, C. M., Leon, M. I., Robin, K. M., & Weinberger, N. M. (2009). Conditioned tone control of brain reward behavior produces highly specific representational gain in the primary auditory cortex. Neurobiology of Learning and Memory, 92(1), 27–34. https://doi.org/https://doi.org/10.1016/j.nlm.2009.02.008

Kelly, J. B., & Masterton, B. (1977). Auditory sensitivity of the albino rat. J Comp Physiol Psychol, 91(4), 930–936. https://doi.org/10.1037/h0077356

Koivuniemi, A., Wilks, S. J., Woolley, A. J., & Otto, K. J. (2011). Chapter 10 - Multimodal, longitudinal assessment of intracortical microstimulation. In J. Schouenborg, M. Garwicz, & N. Danielsen (Eds.), Progress in Brain Research (Vol. 194, pp. 131–144). Elsevier. https://doi.org/https://doi.org/10.1016/B978-0-444-53815-4.00011-X

Koivuniemi, A. S., Regele, O. B., Brenner, J. H., & Otto, K. J. (2011). Rat behavioral model for high-throughput parametric studies of intracortical microstimulation. Annu Int Conf IEEE Eng Med Biol Soc, 2011, 7541–7544. https://doi.org/10.1109/iembs.2011.6091859

Kozai, T. D. Y., Marzullo, T. C., Hooi, F., Langhals, N. B., Majewska, A. K., Brown, E. B., & Kipke, D. R. (2010). Reduction of neurovascular damage resulting from microelectrode insertion into the cerebral cortex using in vivo two-photon mapping. Journal of Neural Engineering, 7(4). https://doi.org/10.1088/1741-2560/7/4/046011

Kramer, D. R., Kellis, S., Barbaro, M., Salas, M. A., Nune, G., Liu, C. Y., Andersen, R. A., & Lee, B. (2019). Technical considerations for generating somatosensation via cortical stimulation in a closed-loop sensory/motor brain-computer interface system in humans. Journal of Clinical Neuroscience, 63. https://doi.org/10.1016/j.jocn.2019.01.027

Kunigk, N. G., Urdaneta, M. E., Malone, I. G., Delgado, F., & Otto, K. J. (2022). Reducing Behavioral Detection Thresholds per Electrode via Synchronous, Spatially-Dependent Intracortical Microstimulation. Frontiers in Neuroscience, 16. https://doi.org/10.3389/fnins.2022.876142

Levitt, H. (1971). Transformed Up-Down Methods in Psychoacoustics. The Journal of the Acoustical Society of America, 49(2B). https://doi.org/10.1121/1.1912375

Liu, J., Earp, J. C., Lertora, J. J. L., & Wang, Y. (2022). Chapter 19 - Dose-effect and concentrationeffect analysis. In S.-M. Huang, J. J. L. Lertora, P. Vicini, & A. J. Atkinson (Eds.), Atkinson’s Principles of Clinical Pharmacology (Fourth Edition) (pp. 359–376). Academic Press. https://doi.org/https://doi.org/10.1016/B978-0-12-819869-8.00039-2

Lycke, R., Kim, R., Zolotavin, P., Montes, J., Sun, Y., Koszeghy, A., Altun, E., Noble, B., Yin, R., He, F., Totah, N., Xie, C., & Luan, L. (2023). Low-threshold, high-resolution, chronically stable intracortical microstimulation by ultraflexible electrodes. bioRxiv. https://doi.org/10.1101/2023.02.20.529295

Macmillan, N. A., & Creelman, C. D. (2005). Detection theory: A user’s guide, 2nd ed. Lawrence Erlbaum Associates Publishers.

Mekarski, J. E. (1988). Main effects of current and pimozide on prepared and learned self-stimulation behaviors are on performance not reward. Pharmacology Biochemistry and Behavior, 31(4), 845–853. https://doi.org/https://doi.org/10.1016/0091-3057(88)90394-2

Müller, H.-G., & Schmitt, T. (1990). Choice of Number of Doses for Maximum Likelihood Estimation of the ED50 for Quantal Dose-Response Data. Biometrics, 46(1), 117–129. https://doi.org/10.2307/2531635

Ni, A. M., & Maunsell, J. H. R. (2010). Microstimulation Reveals Limits in Detecting Different Signals from a Local Cortical Region. Current Biology, 20(9), 824–828. https://doi.org/10.1016/j.cub.2010.02.065

Öztürk, S., Devecioğlu, I., Beygi, M., Atasoy, A., Mutlu, Ş., Özkan, M., & Güçlü, B. (2019). Real-Time Performance of a Tactile Neuroprosthesis on Awake Behaving Rats. IEEE Transactions on Neural Systems and Rehabilitation Engineering, 27(5), 1053–1062. https://doi.org/10.1109/TNSRE.2019.2910320

Page, D. M., George, J. A., Wendelken, S. M., Davis, T. S., Kluger, D. T., Hutchinson, D. T., & Clark, G. A. (2021). Discriminability of multiple cutaneous and proprioceptive hand percepts evoked by intraneural stimulation with Utah slanted electrode arrays in human amputees. Journal of NeuroEngineering and Rehabilitation, 18(1), 12. https://doi.org/10.1186/s12984-021-00808-4

Pancrazio, J. J., Deku, F., Ghazavi, A., Stiller, A. M., Rihani, R., Frewin, C. L., Varner, V. D., Gardner, T. J., & Cogan, S. F. (2017). Thinking Small: Progress on Microscale Neurostimulation Technology. Neuromodulation, 20. https://doi.org/10.1111/ner.12716

Pankevich, D. E. (2012). Animals in Neuroscience Research. In I. o. M. U. N. R. C. (US) (Ed.), International Animal Research Regulations: Impact on Neuroscience Research: Workshop Summary. Washington (DC): National Academies Press (US). https://www.ncbi.nlm.nih.gov/books/NBK100126/

Potter, K. A., Buck, A. C., Self, W. K., & Capadona, J. R. (2012). Stab injury and device implantation within the brain results in inversely multiphasic neuroinflammatory and neurodegenerative responses. Journal of Neural Engineering, 9(4), 046020. https://doi.org/10.1088/1741-2560/9/4/046020

Rajan, A. T., Boback, J. L., Dammann, J. F., Tenore, F. V., Wester, B. A., Otto, K. J., Gaunt, R. A., & Bensmaia, S. J. (2015). The effects of chronic intracortical microstimulation on neural tissue and fine motor behavior. Journal of Neural Engineering, 12(6), 066018. https://doi.org/10.1088/1741-2560/12/6/066018

Riley, J. R., Borland, M. S., Tamaoki, Y., Skipton, S. K., & Engineer, C. T. (2021). Auditory Brainstem Responses Predict Behavioral Deficits in Rats with Varying Levels of Noise-Induced Hearing Loss. Neuroscience, 477, 63–75. https://doi.org/https://doi.org/10.1016/j.neuroscience.2021.10.003

Rousche, P. J., & Normann, R. A. (1999). Chronic intracortical microstimulation (ICMS) of cat sensory cortex using the utah intracortical electrode array. IEEE Transactions on Rehabilitation Engineering, 7(1). https://doi.org/10.1109/86.750552

Schindler, C. W., Thorndike, E. B., & Goldberg, S. R. (1993). Acquisition of a nose-poke response in rats as an operant. Bulletin of the Psychonomic Society, 31(4), 291–294. https://doi.org/10.3758/BF03334932

Schmidt, E. M., Bak, M. J., Hambrecht, F. T., Kufta, C. V., O’Rourke, D. K., & Vallabhanath, P. (1996). Feasibility of a visual prosthesis for the blind based on intracortical micro stimulation of the visual cortex. Brain, 119(2), 507–522. https://doi.org/10.1093/brain/119.2.507

Shannon, R. V. (1992). A Model of Safe Levels for Electrical Stimulation. IEEE Transactions on Biomedical Engineering, 39(4). https://doi.org/10.1109/10.126616

Sloan, A. M., Dodd, O. T., & Rennaker, R. L. (2009). Frequency discrimination in rats measured with tone-step stimuli and discrete pure tones. Hearing Research, 251(1), 60–69. https://doi.org/https://doi.org/10.1016/j.heares.2009.02.009

Sturgill, B., Radhakrishna, R., Thai, T. T. D., Patnaik, S. S., Capadona, J. R., & Pancrazio, J. J. (2022). Characterization of Active Electrode Yield for Intracortical Arrays: Awake versus Anesthesia. Micromachines, 13(3). https://doi.org/10.3390/mi13030480

Tehovnik, E. J. (1996). Electrical stimulation of neural tissue to evoke behavioral responses. Journal of Neuroscience Methods, 65. https://doi.org/10.1016/0165-0270(95)00131-X

Urdaneta, M. E., Kunigk, N. G., Currlin, S., Delgado, F., Fried, S. I., & Otto, K. J. (2022). The Long-Term Stability of Intracortical Microstimulation and the Foreign Body Response Are Layer Dependent. Frontiers in Neuroscience, 16. https://doi.org/10.3389/fnins.2022.908858

Urdaneta, M. E., Kunigk, N. G., Delgado, F., Fried, S. I., & Otto, K. J. (2021). Layer-specific parameters of intracortical microstimulation of the somatosensory cortex. Journal of Neural Engineering, 18(5). https://doi.org/10.1088/1741-2552/abedde

Vasilev, D., Havel, D., Liebscher, S., Slesiona-Kuenzel, S., Logothetis, N. K., Schenke-Layland, K., & Totah, N. K. (2021). Three Water Restriction Schedules Used in Rodent Behavioral Tasks Transiently Impair Growth and Differentially Evoke a Stress Hormone Response without Causing Dehydration. eneuro, 8(6), ENEURO.0424-0421.2021. https://doi.org/10.1523/ENEURO.0424-21.2021

